# Characterization of Somatic Mutations in Human tRNA Genes Reveals Tumor-Specific Mutational Loads, and the Generation of tRNA Variants that Alter the Genetic Code

**DOI:** 10.1101/2025.06.09.658696

**Authors:** Marina Murillo-Recio, Marina Salvadores, Adrián Gabriel Torres, Lina Tsapanou, Aina Vaquer Picó, Fran Supek, Lluís Ribas de Pouplana

## Abstract

Transfer RNA genes (tDNAs) are essential genomic elements that safeguard translational fidelity. Using the T2T version of the human genome we have mapped the position of human tDNAs and analyzed their individual transcriptional activities. Then we have characterized, at single base resolution, the impact of somatic mutations in human tDNAs and its relationship to the transcriptional status of each gene. We confirm that tDNAs are hotspots for somatic mutagenesis, and show that they display mutational loads that are directly proportional to their transcription rates. Highly transcribed tDNAs in tumors or healthy tissues accumulate mutations at rates up to nine-fold higher than highly transcribed protein-coding genes. Mutational loads at tDNAs are tumor-specific, and increase with patient age. Mutations at structurally conserved tRNA positions appear to be under negative selection. Strikingly, anticodon nucleotides crucial for decoding frequently acquire somatic mutations, readily generating chimeric tRNAs species capable of systematically introducing amino acid substitutions across the proteome. Our results reveal a previously unrecognized source of somatic heterogeneity in human cancer and aging tissues that may directly impact upon translation efficiency and fidelity, and cause cell-specific proteostasis degeneration.

## Introduction

The translational machinery of cells is finely tuned to maintain proteostasis, a key factor in cellular survival and proliferation (Gingold et al. 2014; Goodarzi et al. 2016; Torrent et al. 2018). Transfer RNAs (tRNAs) are central to translation, pairing their anticodons to codons in messenger RNAs and delivering the corresponding amino acids during translation. The human genome encodes over 600 tRNA genes (tDNAs) (Chan and Lowe 2016), which are transcribed by RNA polymerase III (Pol III) via the recognition of two internal promoter regions that are highly conserved among all tDNAs (Fig. 1a) (Galli et al. 1981; Giege et al. 1998). All active tRNAs share a common three-dimensional structure required for their function in the ribosome, but each tRNA must present idiosyncratic recognition motifs essential to preserve the faithful translation of the Genetic Code (Fig. 1a) (Giege et al. 1998; Berg and Brandl 2021), which takes place when tRNAs are aminoacylated at their 3’ end by aminoacyl-tRNA synthetases (ARS) (Giege et al. 1998; Ibba and Soll 2000). The identity elements used for accurate aminoacylation are idiosyncratic to each tRNA-ARS cognate pair and, in most cases, include the bases at the anticodon (Giege et al. 1998; Giege and Eriani 2023). However, some ARS do not interact with the anticodon sequence of their cognate tRNAs, and mutations at these anticodons can lead to chimeric tRNAs that cause amino acid substitutions throughout the proteome (Geslain et al. 2010; Parisien et al. 2013; Reverendo et al. 2014; Santos et al. 2018; Pinzaru and Tavazoie 2023).

**Figure 1.**
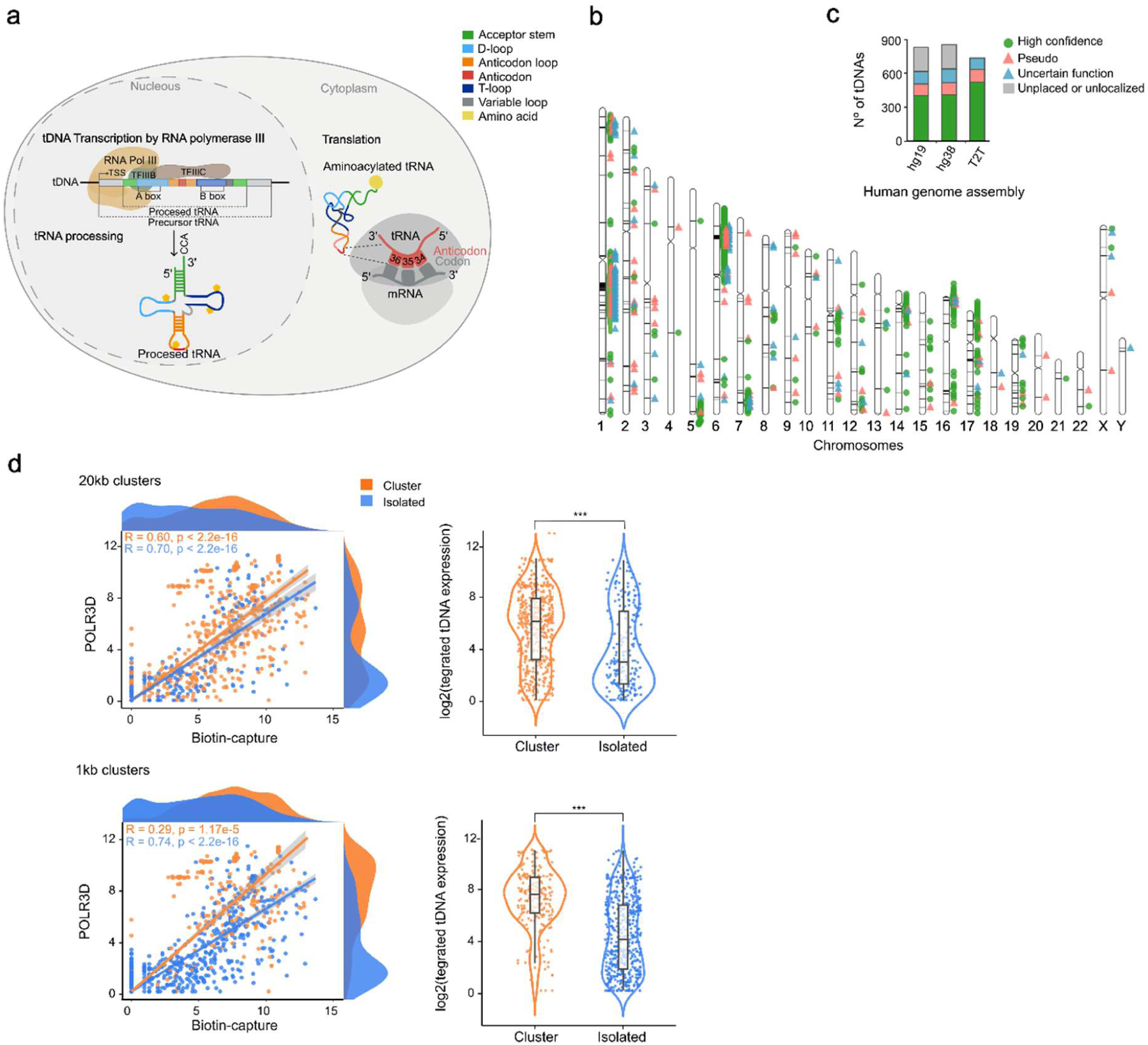
tDNAs localization and transcription. **(a)** Schematic representation of tDNA biology including tDNA transcription by Pol III, tRNA processing, tRNA structure and codon-anticodon pairing. The diagram illustrates the conserved internal promoter regions (A box and B box), which are essential for RNA Polymerase III-mediated transcription. These regions facilitate the recruitment of transcription factors TFIIIB and TFIIIC. The precursor tRNA undergoes processing steps, including the addition of a 3′ CCA tail, necessary for amino acid attachment through aminoacylation by aminoacyl-tRNA synthetase, ultimately producing a fully mature and functional tRNA. In the cytoplasm, mature tRNA participates in protein synthesis by base-pairing the anticodon (positions 34 to 36) with complementary mRNA codons and delivering the appropriate amino acid. **(b)** Ideogram showing the chromosomal distribution and localization of human tDNAs in the T2T genome assembly including tRNA-scan-SE 2.0 prediction classification (high-confidence, pseudogenes, and tDNAs with uncertain function). **(c)** Quantification and classification of tDNAs across different human genome assemblies (hg19, hg38, T2T) based on analyses conducted using tRNAscan-SE 2.0. tDNAs identified within unplaced or delocalized sequences of the reference genome are in grey. **(d)** Correlation between POLR3D and biotin-capture expression values by tDNA localization (cluster or isolated). tDNAs that are not in clusters are identified as isolated tDNAs. On the top row: Analysis based on clusters within 20 kb. Bottom row: Analysis based on clusters within 1 kb. On the left scatter plots showing the relationship between POLR3D and biotin capture expression values for tDNAs classified as either clustered (orange) or isolated (blue). The plots include Spearman correlation (R) and the associated *p* values. On the right, the violin plots show the log2 integrated tDNA expression for clustered and isolated tDNAs, with Wilcoxon significance levels indicated as ***, p ≤ 0.001. For the 20 kb clusters, p < 2.2e-16, and for the 1 kb clusters, p = 1.714e-10.

Alterations in tRNA populations are linked to numerous human diseases (Orellana et al. 2022). For example, increased tRNA abundance is a feature of cancer cells, which adapt their tRNA populations to optimize the translation of specific genetic programs (Goodarzi et al. 2016; Zhang et al. 2018; Gupta et al. 2022; Pinzaru and Tavazoie 2023). Quantitative and qualitative alterations in tRNA populations -including tRNA-derived fragments (tRFs)- are widespread in human cancers (Huang et al. 2018; Cabrelle et al. 2024),where they have been linked to aberrant signaling and translational reprogramming in several cancer types, including lung and breast cancer (Skeparnias et al. 2020; Kwon et al. 2021). Specific tRNA isoacceptors and post-transcriptional base modifications regulate cancer cell survival and influence metastatic potential by controlling the translation of proliferation-related genes (Earnest-Noble et al. 2022; Garcia-Vilchez et al. 2023). Moreover, metabolic and therapeutic stresses, such as chemotherapy, induce codon-biased aberrant protein production through altered tRNA function and decoding. Thus, variations in tRNA populations reshape the cancer proteome and support tumor adaptation and therapy resistance (Kochavi et al. 2024; Wernaart et al. 2024; Yang et al. 2024).

Somatic mutagenesis causes changes in the DNA sequence of somatic cells, drives the emergence of cancers, and is an important contributor to the aging process (Vijg and Dong 2020). The nature and dynamics of human somatic mutagenesis has been studied mostly in protein-coding genes, and its impact upon non-coding RNA genes is less understood. The activity of members of the APOBEC3 subfamily of cytidine deaminase enzymes is an important contributor to somatic mutagenesis (Taylor et al. 2013; Mas-Ponte and Supek 2020; Langenbucher et al. 2021; Sanchez et al. 2023). APOBEC3 enzymes function, primarily, as an innate immunity mechanism to defend against viruses and mobile genetic elements but, in some cancers, APOBEC3A and APOBEC3B paralogs can generate a high mutational burden (Burns et al. 2013; Roberts et al. 2013; Nik-Zainal et al. 2014; Supek and Lehner 2017; Petljak et al. 2022). APOBECs can access only single-stranded nucleic acids, and are known to act upon ssDNA intermediates during DNA repair, as well as ssDNA segments within structured, stem-loop DNA (Roberts et al. 2013; Buisson et al. 2019; Mas-Ponte and Supek 2020).

Studies in bacteria and yeast have reported that non-coding genes can accumulate mutations also caused by the activity of APOBEC3 enzymes. In an *E. coli* model, mutational impact of heterologous expressed APOBEC3B is highest in genes transcribed by Pol III, such as tDNAs (Sakhtemani et al. 2019). Mutational studies in a yeast model also suggested that human APOBEC3B overexpression causes severe DNA damage in Pol III-transcribed genes (Saini et al. 2017), suggesting a transcription-related mechanism. In human germline cells, mutations in nuclear tDNAs are subject to strong purifying selection (Seplyarskiy et al. 2023), (Thornlow et al. 2018), but analysis of tRNA sequences obtained from the plasma of healthy human volunteers revealed significant levels of human heterogeneity (Parisien et al. 2013; Berg et al. 2019). Human tDNAs are known to accumulate somatic mutations at high rates, and APOBEC has been identified as a major contributor to this phenomenon (Sakhtemani et al. 2019). However, a detailed analysis of tDNA somatic mutations, including their distribution at single-base resolution and their potential physiological impact is lacking.

We have mapped all tDNAs in the T2T version of the human genome (Fig. 1b), and used this information to study somatic mutagenesis at these loci (Hoyt et al. 2022; Nurk et al. 2022). We identify 733 tDNA genes (more than 100 than previously thought), leaving no unplaced or unlocalized genes (Fig. 1c). As previously reported, tDNAs can be found isolated or in large gene clusters (Bermudez-Santana et al. 2010; Van Bortle et al. 2017). Combining our data with POLR3D ChIP-seq and ‘biotin-capture of nascent RNA’ data, we find that both isolated and clustered tRNAs can be actively transcribed,but tDNAs in clusters exhibit higher activity levels than isolated genes (Fig. 1d).

We then characterized in detail the somatic mutations affecting tDNAs in human tissues, which are prevalent both in tumors and in healthy tissues. Indeed, tDNAs constitute hotspots of somatic mutagenesis whose intensity is directly linked to the transcriptional activity of each gene. Mutational load at tDNAs accumulate at rates up to nine-fold higher than in protein coding genes, and are higher than in other genes transcribed by Pol III. Mutational loads at tDNAs vary greatly between tumor types, display mutational signatures that are different in cancer or in healthy tissues, and readily accumulate with age. Nucleotides important for tRNA structure accumulate mutations at rates lower than expected, but all three anticodon positions have average mutational frequencies. Somatic mutations at anticodon bases at highly transcribed tDNAs may result in mutant chimeric tRNAs capable of introducing specific and ubiquitous amino acid substitutions throughout the proteome.

## Results

### Genome-wide identification of tDNAs and their transcriptional activity

To obtain an accurate picture of the set of human tDNAs we first mapped the positions of these genes in the most recent version of the human genome assembly Jan. 2022 (T2T-CHM13 v2.0/hs1) (Hoyt et al. 2022) (Fig. 1b). Our analysis sets the total number of identified tDNAs to 733, of which 521 tDNAs are confidently predicted as functional tRNAs (∼112 more genes than previously reported with older assemblies) (Fig. 1c). We successfully localize all known tDNAs and remove previous uncertainties regarding the localization of some of these genes (Fig. 1c). As previously reported, tDNAs can be found as isolated genes, or as part of large gene clusters that are present in several chromosomes (Bermudez-Santana et al. 2010; Van Bortle et al. 2017) (Fig. 1b). To better characterize the tDNAs, we calculated their genomic distances (Supp. Fig. 1a) and grouped them into clusters based on thresholds of 20 kb or 1 kb (Supp. Fig. 1b and Supp. Table. 1). The size of the resulting clusters varies significantly: 1 kb clusters typically contain 2–5 tDNA genes, while 20 kb clusters can range from 2 to 112 tDNAs (Supp. Fig. 1c). The distribution and contents of clusters of tDNAs are strongly conserved among primates, with a tendency to larger tDNA clusters in Homo sapiens (Bermudez-Santana et al. 2010) (Supp. Fig. 1d).

tDNA transcriptional activity can be estimated by combining POLR3D ChIP-seq data with biotin-capture of nascent RNA (Supp. Table. 2) (Van Bortle et al. 2017). Through this approach, we discovered that while both isolated and clustered tRNAs can be actively transcribed, tDNAs in clusters exhibit higher activity levels than isolated genes (see Fig. 1d) (tDNAs with higher levels of expression are located in clusters). This conclusion holds true even when sets of tRNA isoacceptors are analyzed individually (Supp. Fig. 1e).

### tDNAs are hotspots of somatic mutagenesis

To quantify the extent of somatic mutagenesis tDNAs we used whole-genome sequencing data (WGS) from 9596 samples of cancerous tissues and 1192 samples of clonal healthy tissues. Variant calling data was previously obtained using the hg19 genome, to confirm tRNA variation we used LiftOver to convert T2T tDNAs coordinates to hg19, and we were able to localize 537 tDNAs (Supp. Table. 3). In contrast to previous studies in germinal cells (Thornlow et al. 2018), we confirm that tDNAs in somatic cells are mutational hotspots in tumor samples (Fig. 2a).

**Figure 2.**
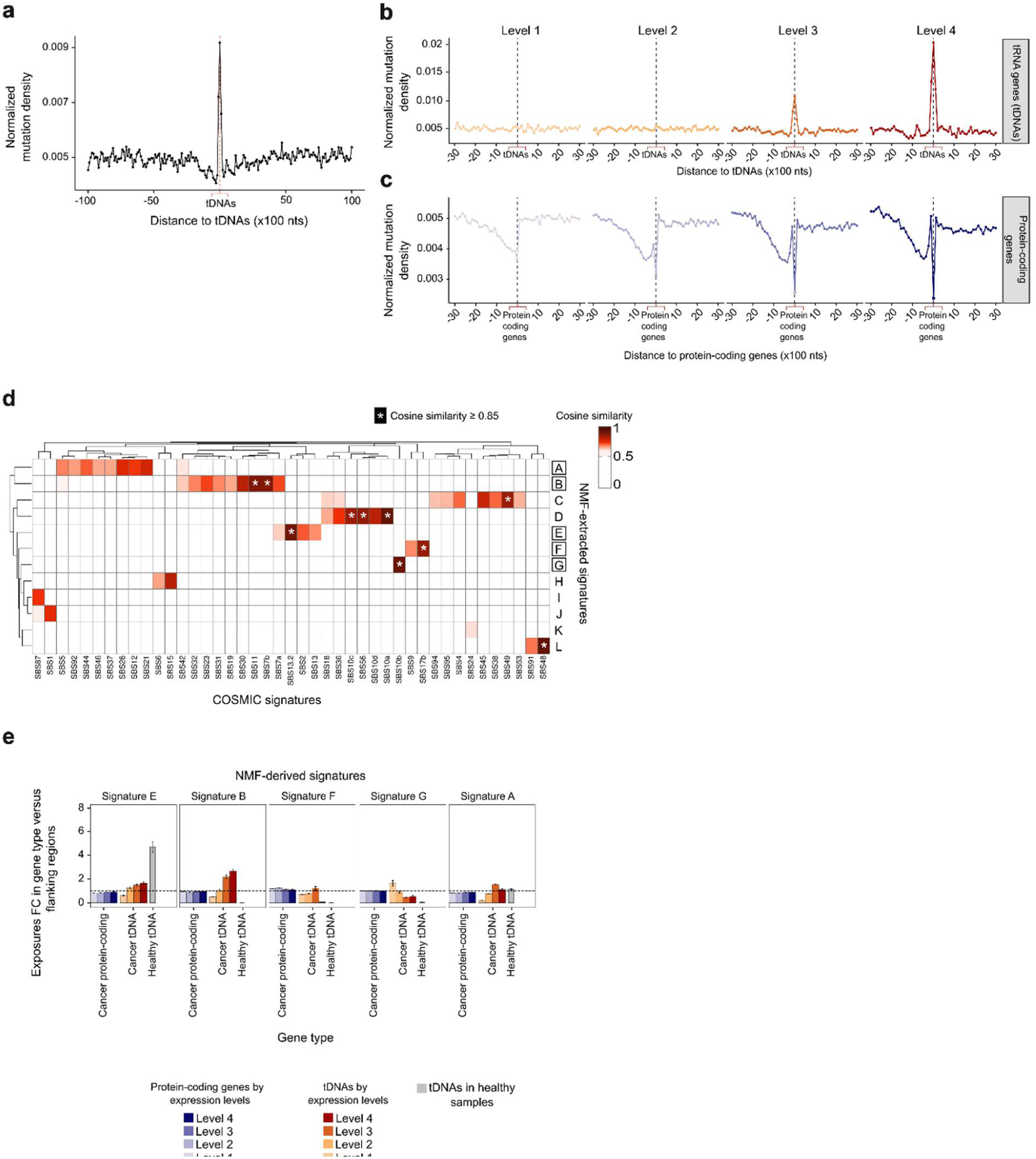
Somatic mutational profile and mechanism in tDNAs. **(a)** Normalized mutation density in tDNAs of cancer samples, including flanking regions (10 kb downstream and upstream divided into windows of 100 nucleotides). **(b)** Normalized mutation density in tDNAs classified by transcription levels from highest (Level 4) to lowest (Level 1), including flanking regions (3 kb upstream and downstream divided into 100 nucleotide windows). **(c)** Normalized mutation density in protein-coding genes classified by expression levels and flanking regions (3 kb upstream and downstream divided into windows of 100 nucleotides). **(d)** Comparison of NMF-extracted signatures with COSMIC signatures. Heatmap showing cosine similarity values between the trinucleotide spectra of our de-novo NMF-derived signatures (rows) and the reference COSMIC signatures (columns). Only COSMIC signatures with a cosine similarity ≥ than 0.5 are shown. A COSMIC signature is considered to be assigned when the cosine similarity is ≥ 0.85, indicated by an asterisk (*). (**e)** Sample exposure ratio (fold-change, FC) between genes (stratified by type into tDNAs or protein-coding genes) and their flanking regions, for each NMF-extracted signature. Exposures are analyzed for tDNAs and protein-coding genes in cancer samples, as well as for tDNAs in healthy samples. Data for both gene types in cancer samples is stratified by expression levels, from low (Level 1) to high (Level 4).

The internal mutational burden of tDNAs correlates directly with their transcriptional activity (Fig. 2b). Silent tDNA genes show no mutation accumulation, while highly transcribed tDNA genes accumulate internal sequence mutations at rates up to nine times higher than in protein-coding genes (Fig. 2c). This difference suggests a direct link between Pol III activity and the mutation accumulation in tDNA genes. While tDNAs transcription is driven by Pol III, protein-coding genes are transcribed by RNA polymerase II (Pol II), which recruits DNA repair mechanisms such as transcription-coupled nucleotide excision repair (TCR) (Guo et al. 2022). In comparison, Pol III-transcription lacks known coupled-repair, lacking mechanisms to deal with the Transcription-Associated Mutagenesis (TAM) phenomena (Jinks-Robertson and Bhagwat 2014; Thornlow et al. 2018). We asked whether the observed tDNA somatic mutation load is also detectable in other Pol III-transcribed genes such as ribosomal RNA (rRNA), small nuclear RNA (snRNA) and unclassified RNAs (miscRNA), as was reported in the human germline mutations (Seplyarskiy et al. 2023). Mutational densities at rRNAs, snRNAs, and miscRNAs are significantly lower than at tDNAs and display different mutational profiles, suggesting that the mechanisms leading to somatic mutations in tDNAs are gene-type specific (Supp. Fig. 2). We quantified the tDNA mutational burden in clonal samples of healthy tissues (including brain, colon, liver, and lung) using whole-genome sequencing (WGS) data, and again found a significant accumulation of somatic mutations in tDNAs (Fig. 3b). Thus, somatic tDNA mutations are also prevalent in healthy tissues, where their time-wise accumulation might contribute to the proteostasis defects characteristic of aging individuals.

**Figure 3.**
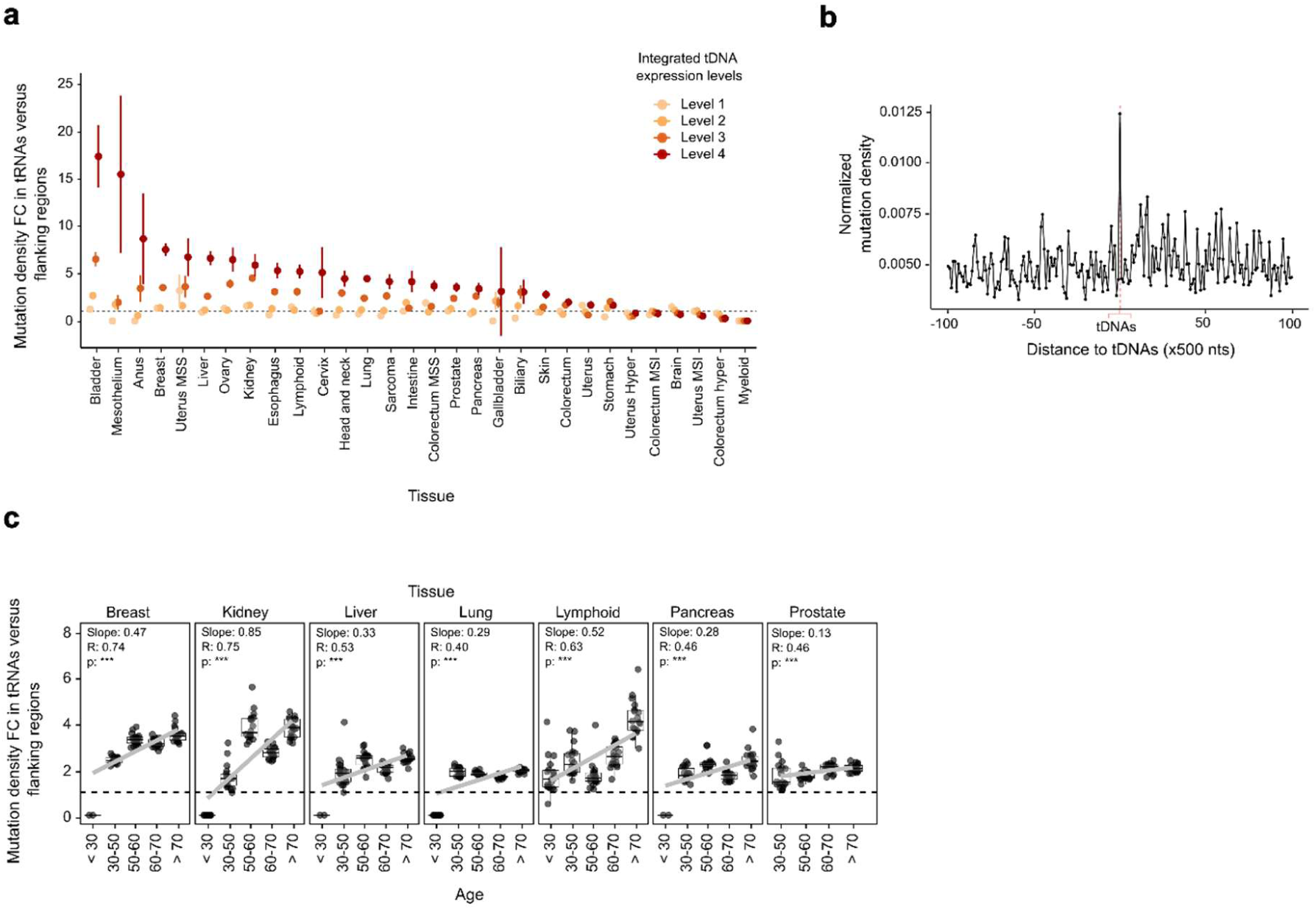
Age-related and tissue-specific mutational burdens in somatic tDNAs. **(a)** Mutation profiles in tDNAs across tissue types (including tissues with n > 20) relative to the flaking regions. Each dot represents the mean FC between mutation density values for tDNAs and the flanking regions (see Methods). tDNAs are categorized based on expression levels. **(b)** Normalized mutation density in tDNAs from healthy non-cancerous tissues (flanking regions of 50 kb downstream and upstream di-vided into windows of 500 nucleotides). **(c)** Tissues showing a positive correlation between muta-tional density in tDNAs and age. Data is grouped by tissue and age range (including tissues with n > 200). Each dot represents the FC between mutational densities of tDNAs and a flanking 10-window set (see Methods). The plots include values for the slope, spearman correlation coefficient (R), and statistical significance of the correlation (***,*p ≤ 0.001*). Results for other tissues Supp. Fig. 6 and detailed statistics are available in.

### Mutational signatures and role of APOBEC3 in tDNA Mutagenesis

The nature and frequency of somatic mutations generate a signature that can be used to identify the agents responsible for the observed mutations, based on the frequencies of trinucleotide contexts of the mutations. Over a hundred such mutational signatures exist in human samples, and mutational agents are identified for approximately half of them (Alexandrov et al. 2020; Degasperi et al. 2022). APOBEC mutations generate a clear signature of C>T and C>G changes restricted to T<u>C</u>W contexts (where W is T or A). Mutational signature analyses performed specifically at tDNA loci detected two main mutational signatures, SBS13 and SBS2, (Fig. 2d and Supp. Fig. 7) (Alexandrov et al. 2020), both identifying APOBEC3-driven mutagenesis as the major contributor to the observed tDNA mutations. The APOBEC3 signature in tRNA genes was also present in genomes of healthy cells (Fig. 2e and Supp. Fig. 4). While genome-wide signatures of APOBEC3 mutagenesis are rare in healthy somatic cells, in contrast to cancer genomes (Franco et al. 2019 Genome Biology), we do observe this mutagenesis signature in tDNAs both in tumors and healthy tissues. Therefore, high transcription rates and/or other factors contributing to ssDNA occurrence must act at tRNA loci to facilitate APOBEC mutagenesis therein. Other genes transcribed by Pol III display different mutational signatures, indicating that APOBEC3-driven mutagenesis at tDNA genes is not driven only by high levels of transcription, but likely requires additional factors specific to tDNAs. We suspected secondary-structure prone DNA sequences, and we used hairpin-localization algorithms to determine the presence of APOBEC3-preferred mutational sites in tDNAs (Roberts et al. 2013; Buisson et al. 2019; Sui et al. 2020; McCann et al. 2023). This analysis revealed a significant enrichment of DNA hairpin formations in tRNA genes compared to the rest of the genome (Chi-squared test p = 0.01578) or to other Pol III-transcribed genes (Chi-squared test p = 0.03653).

Interestingly, we find that the mutational profiles of tDNAs in healthy tissues and tumors are not identical (Fig. 2e). Some mutational signatures identified in human tumors were in fact negatively correlated with tDNA transcriptional intensity. This suggests a protective effect of active, open chromatin at highly-transcribed tDNAs, possibly through recruiting DNA repair (Polak, Lawrence et al. 2014 Nat Biotechnol). Based on previously reported experiments, base excision DNA repair may be enriched in this situation (Saini, Roberts et al. 2017 DNA Repair). For example, signature F (likely corresponding to COSMIC catalog SBS17b), which is associated to oxidative damage to the free nucleotide pool and also to 5-fluorouracil treatment (Alexandrov et al. 2020), shows decreased incidence in highly-transcribed tDNA genes, where its exposure levels are 8.75 times lower than in tDNAs with low transcription rates (Fig. 2e).

On the other hand, mutational signatures that are prevalent in healthy tissues but not detected in tumor genomes resemble a known signature, SBS10, associated to ultramutation associated to defects of the proofreading domain of the DNA polymerase epsilon (PolE). Given that PolE-associated ultramutagenesis is a known feature of cancers, and expected to be largely absent in healthy cells, our data suggests that this signature originates from another, currently unclear cause. Speculatively, factors such as transcription-replication conflicts, or usage of replication origins which can be collocated with tDNAs (noted in yeast), could be altered in cancer, resulting in a tumor-specific mutational process at tDNA.

### Tumor type- and age-dependence of tDNA somatic mutagenic rates

The overexpression of tDNAs in human cancers has been repeatedly reported (Goodarzi et al. 2016; Zhang et al. 2018; Gupta et al. 2022; Orellana et al. 2022; Pinzaru and Tavazoie 2023), and linked to adaptive mechanisms of codon-anticodon optimization that prioritize the translation of genetic programs important for tumor growth (Goodarzi et al. 2016). Moreover, tDNA expression can be cell type- or tissue-specific, often resulting in differential expression of not only mature tRNAs but also of tRNA fragments (Dittmar et al. 2006; Torres et al. 2019). To test whether somatic tDNA mutagenesis is a distinguishing feature of different human tissues we compared tDNA somatic mutagenesis by cancer type. This analysis revealed that tDNA mutational load varies by cancer type and tissue, with bladder cancer (BLCA) exhibiting the highest rates of tDNA mutagenesis (Fig. 3a). In agreement with our previous finding that high levels of transcription correlate with mutational frequencies at tDNAs, cancer types characterized by up-regulated levels of tRNA expression (including bladder, uterine corpus endometrial carcinoma and breast cancer)(Zhang et al. 2018), also exhibit the highest mutational burdens in tDNAs.

Next, we investigated whether mutational loads in tDNAs accumulate with age by comparing samples from cancer patients across different age groups. Our results demonstrate a significant age-related increase in tDNA somatic mutational load in various cancer types, particularly in tumors that display high levels of tDNAs transcription. These results suggest that the accumulation of somatic mutations in tDNAs is a characteristic of aging human tissues (Fig. 3c and Supp. Fig. 6). tDNAs from donors below thirty years of age show no signs of mutational load, while these become conspicuous in the forty-to-sixty age range, and increase further in older patients. Available data from healthy human tissues is insufficient to determine the age-related distribution of somatic mutations at tDNAs, but the behavior observed in tumors, and the fact that the datasets from healthy tissues do reveal high levels of somatic mutations at tDNAs, indicate that these mutations do accumulate in cells as humans age.

### Distribution of somatic mutations in tDNAs and evidence for negative selection

While genomic signatures of negative selection in somatic cells are quite subtle, some protein-coding genes, including essential genes and oncogenes do display lower mutation rates than expected (Van den Eynden et al. 2016; Martincorena et al. 2017; Besedina and Supek 2024). We reasoned that the increased rate of somatic mutagenesis at tDNA loci, coupled with the fact that hundreds of tDNA gene copies can be easily aligned to compare variation at individual nucleotides, might provide good statistical power to search for evidence of negative selection acting against somatic mutations at critical positions of the tRNA molecule.

Thus, we quantified tDNA mutational rates at single base resolution for all human tDNAs to search for negative selection acting upon mutations at specific sites. This analysis revealed that most universally conserved positions in tRNA sequences (Beuning and Musier-Forsyth 1999) display mutation rates significantly lower than the rest of nucleotides in tRNAs (Fig. 4a, Fig. 4b ad Supp. Fig. 7), indicating that these positions experience negative selection in somatic cells that eliminates mutations at these sites. Interestingly, this analysis revealed other positions that, while not universally conserved, also display lower than expected mutagenic rates, suggesting that these nucleotides play important, but as yet unidentified, functional roles in tRNAs (Fig. 4a and Supp. Fig. 7). Possibly, mutations at these positions generate aberrant tRNA species that directly interfere with the translation machinery and significantly affect cell fitness.

**Figure 4.**
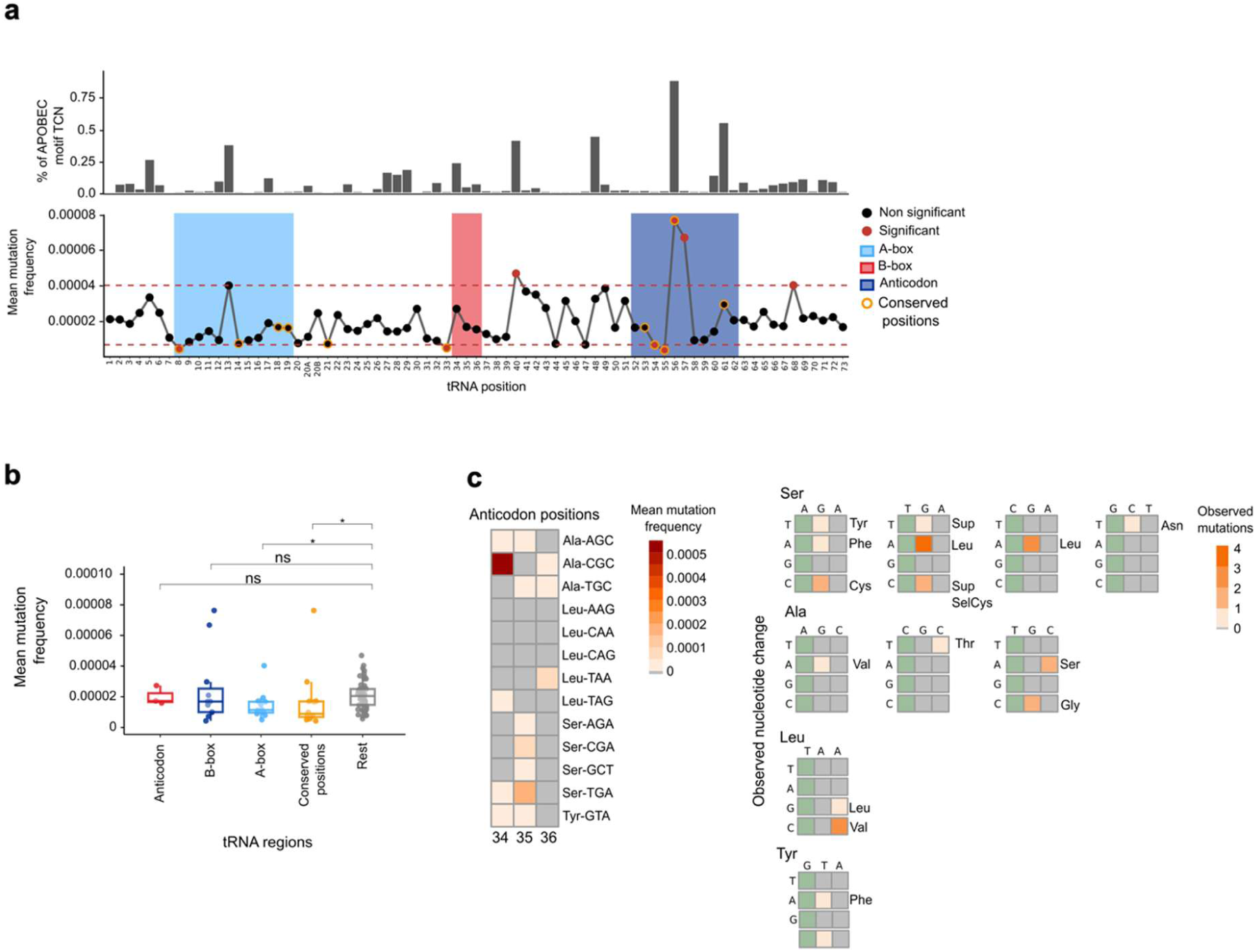
Somatic tDNA mutation frequencies at single-base resolution and distribution of APOBEC motif presence across tDNAs genomic positions. **(a)** The lower panel shows mutation rates at single-base resolution across tRNA sequences, highlighting universally conserved positions (U8, A14, G18, G19, A21, U33, G53, T54, Ψ55, C56, A58, C61, C74, C75), internal promoter regions such as the A-box (positions 8–19), the B-box (positions 52–62), and the anticodon (positions 34–36). Statistically significant positions are indicated with red dots. Specific tDNA positions are considered significant when adj. *p ≤* 0.01 and if its mean mutation density exceeds the 0.95 quantile or falls below the 0.5 quantile of the overall distribution (quantile thresholds are indicated by red dashed lines). The upper panel shows the percentage of tDNAs with APOBEC-associated TCN motifs (where N = T, C, G, or A) across tDNA gene sequences. Internal hotspots for somatic mutations closely match the distribution of TCN signatures in tDNA genes. **(b)** Comparison of mutation densities in described functionally and structurally important tRNA positions versus other positions. Wilcoxon significance levels are indicated in the plot: A-box vs. rest (adj. *p* = 0.025, *), conserved positions vs. rest (adj. *p* = 0.025, *), anticodon vs rest (adj. *p* = 0.971, ns), and B-box vs rest (adj. *p* = 0.541, ns). **(c)** Characterization of mutational profiles in the anticodon region of potential chimeric tRNAs, specifically for isoacceptors of alanine (Ala), serine (Ser), leucine (Leu), and tyrosine (Tyr). The heatmap on the left shows the mean mutation density at each anticodon position, classified by isodecoders for Ala, Ser, Leu and Tyr. The heatmap on the right illustrates the number of times specific mutational changes were observed in the anticodon region for specific isodecoders of Ala, Ser, Leu and Tyr. Position 34 is highlighted in green, as mutations at this position are always synonymous. Next to each mutation (observed nucleotide change), the codon recognized by the resulting chimeric tRNA is indicated. For example, the tRNA Ala-AGC mutates to Ala-AAC, resulting in a tRNA that still transports alanine but now binds the codon for valine.

Strikingly, positions critical for translation elongation such as the anticodon triplet, where mutations can severely disrupt the efficiency and fidelity of protein synthesis (Geslain et al. 2010; Reverendo et al. 2014; Santos et al. 2018; Vijg and Dong 2020; Pinzaru and Tavazoie 2023), showed no evidence of decreased mutation rates (Fig. 4a and Fig. 4b), and we identified several instances of mutations at anticodon positions (see below).

In order to analyze the regional mutation load in structural domains of the tRNA, or at the two internal promoters of tDNAs, we used a rolling windows method to calculate the relative mutation load in ten-nucleotide sections of these molecules. This revealed that tDNA genes experience higher mutational loads at their 3’ halves (Supp. Fig. 8). Interestingly, the second promotor region (B-box) is where most mutations accumulate (Fig. 4a and Fig. 4b). This mutational distribution might reflect a locally protective effect of the Pol III complex of the transcription factor TFIIIB over a section of tDNA positive strands during their transcription. Alternatively, the presence of sequences or motifs required for the activity of APOBEC in the 3’ half of tDNAs could be responsible for the relative accumulation of mutations in these regions of tDNAs. Consistent with this second possibility, we find that position 56, within the region of the B box, is the highest mutated position in all tDNAs and is occupied by a highly conserved cytosine in all human tRNAs (Fig. 4a).

### Somatic mutations at tDNAs generate misincorporating tRNAs

The attachment of amino acids to their cognate tRNA molecules by aminoacyl-tRNA synthetases (ARS) is a crucial step for accurate protein translation. For the majority of ARS, tRNA recognition requires interactions with both the acceptor stem and anticodon loop motifs (Hou and Schimmel 1988; Park and Schimmel 1988; Perret et al. 1990; Giege et al. 1998; Rubio Gomez and Ibba 2020; Giege and Eriani 2023). However, ARS cognate for amino acids alanine (Ala), leucine (Leu), serine (Ser), and tyrosine (Tyr) recognize only the acceptor stem of their cognate tRNA substrates (Giege and Eriani 2023). In these cases, somatic mutations at tRNA anticodons can lead to ‘chimeric tRNAs’ whose anticodon triplet will recognize codons that do not correspond to the amino acid carried by the chimeric tRNA. Such molecules lead to translation errors throughout the proteome, caused by the misincorporation of Ala, Ser, Leu, or Tyr at non-cognate codons within the ribosome. Strikingly, in the datasets used in this study we could identify numerous examples of somatic mutations leading to the generation of chimeric tRNAs previously shown to induce proteome-wide amino acid substitutions (Fig. 4c) (Geslain et al. 2010; Santos et al. 2018; Pinzaru and Tavazoie 2023). For example, we detected several examples of mutations at position 35 of tRNA^Ser^(UCA) that result in leucine to serine substitutions throughout the proteome (Fig. 4c). The detection of mutations that compromise the fidelity of the Genetic Code at intensely transcribed tDNAs suggests that aging human cells produce mutagenic tRNAs capable of introducing widespread amino acid changes throughout the proteome, and contribute to widespread proteome mosaicism in tissues.

## Methods

### tDNAs prediction

Reference human genomes were obtained from the UCSC Genome Browser. The assemblies analyzed included Jan. 2022 (T2T-CHM13 v2.0/hs1), Dec. 2013 (GRCh38/hg38), and Feb. 2009 (GRCh37/hg19). tDNAs for each assembly were predicted and annotated using tRNAscan-SE 2.0 (v2.0.9, July 2021) (Chan et al. 2021), with parameters set to search for eukaryotic tRNAs and to display both primary and secondary structure components in the covariance model bit scores to distinguish functional tRNAs from potential pseudogenes. For the hg38 and hg19 assemblies, tDNAs that map to unplaced sequences (DNA fragments associated with a specific chromosome but whose order or orientation cannot be determined) or unassigned sequences (DNA fragments not assigned to any chromosome) were identified and designated as delocalized tDNAs. Subsequently, for the rest of the tDNAs the EukHighConfidenceFilter script from tRNAscan-SE 2.0 was used to assess the fidelity of tDNA predictions. This step classified the predicted tDNAs into several categories: ‘high confidence’ (predictions that exhibit strong sequence and structural alignment with known active tRNA profiles), pseudogenes (which encode inactive products due to sequence alterations like deletions), and ‘uncertain function’ (genes whose function or identity is not clearly established, possibly due to non-canonical structures or sequence deviations from known functional tRNAs). The identified tDNAs in the T2T genome, determined using tRNAscan-SE, are available in Supp. Table 1.

### tDNAs genomic distribution and clusters definition

tDNAs annotations from the most updated human genome (T2T-CHM13 v2.0/hs1) were used to analyze tDNA distribution. In order to visualize the genomic coordinates of tDNAs we used the RIdeogram v0.2.2 package from R. Genomic distances between consecutive tDNAs were computed and used to plot their cumulative distribution using ecdf (empirical cumulative distribution function) (Supp. Fig. 1a). Taking into account the profile obtained from ecdf and previous definitions of tDNA clusters (Bermudez-Santana et al. 2010; Van Bortle et al. 2017), tDNA clusters were defined as groups of consecutive tDNAs separated by a specific genomic distance of either 1 kb (“1 kb cluster”) or 20 kb (“20 kb cluster”). All other tDNAs were categorized as isolated (Supp. Fig. 1b and Supp. Table. 1). The correlation between POLR3D and biotin-capture was assessed using the Spearman correlation coefficient. Statistical differences in log2 integrated tDNA expression levels between clustered and isolated tDNAs were evaluated using the Wilcoxon test.

### Synteny analysis

Reference genomes for human and chimpanzee were obtained from the UCSC Genome Browser. For the human genome T2T-CHM13 v2.0/hs1 Jan. 2022. For the chimpanzee genomeClint_PTRv2/panTro6 Jan. 2018. The reference Lemur (*Microcebus murinus*) genome downloaded from NCBI in Feb. 2017 RefSeq (GCF_000165445.2/Mmur_3.0). To perform synteny analysis, we obtained GTF gene annotation files and DNA sequences for each species from Ensembl (https://www.ensembl.org/Homo_sapiens/Info/Index). These files were modified by removing existing tDNA annotations and incorporating data for 20 kb tDNA clusters, including the start and end coordinates of each cluster and their full nucleotide sequences for each specie. MCScanX (Wang et al. 2012) (https://github.com/tanghaibao/jcvi/wiki/Mcscan-(python-version)(python utility JCVIv0.0.26 (Tang et al. 2024)) was used to perform synteny and collinearity analyses. Macrosynteny analyses were used to compare genomic and cluster compositions. The results were visualized using the karyotype format provided by MCScanX.

### tDNAs expression data

Sequencing data for RNA polymerase III occupancy (POLR3D ChIP-seq) and nascent transcription profiling (biotin-capture) assays were obtained from the Gene Expression Omnibus (GEO) database and are accessible through GEO Series accession number GSE96800 (Van Bortle et al. 2017). Specifically, the subset of samples used for this study came from the monocyte dataset and correspond to the following GEO Sample accession numbers: POLR3D (GSM2544232, GSM2544233) and biotin-capture (GSM2544240, GSM2544241). Following the methodology outlined by Van Bortle et al. (2017), preprocessing of the sequencing data involved trimming low-quality bases and removing adapter sequences using cutadapt v4.1. The processed reads were aligned to the complete human reference genome T2T-CHM13 to minimize false positives, using Bowtie2 v2.4.2 in paired-end and local mode with allowance for one mismatch in the seed (-N 1). Given the repetitive nature of tRNA genes, sequencing reads frequently map to multiple genomic sites, complicating the accurate quantification of tDNA transcription levels. To overcome this, multi-mapping reads were allocated to a single “best” alignment position. This method helps reduce the risk of overestimating tDNA transcription levels that could result from counting multiple alignments, while also preventing potential underestimation by excluding all multi-mapping reads (Van Bortle et al. 2017). Gene counts were obtained using featureCounts from subread v2.0.1. In our analysis, gene annotation files in GTF format, derived from tDNA coordinates predicted by tRNAscan-SE 2.0 were used to report tDNA counts. To simulate precursor tDNA sequences, tDNA coordinates were extended by adding 50 bases both upstream and downstream. Subsequently, gene counts normalization was performed using the estimateSizeFactors function from the DESeq2 package. The log 2 mean between the biological replicates was obtained for each experimental data POLR3D and biotin-capture. Finally integrated tDNA expression values were obtained by computing the average between POLR3D and biotin-capture results (Expression data available in Supp. Table. 2). tDNAs were categorized into four expression levels based on quartiles of integrated tDNA expression, with level 4 representing the highest expression values. This categorization was used to study the relationship between tDNA transcription rates and mutagenesis.

### WGS datasets for somatic mutations

For cancer samples, somatic single nucleotide variants (SNVs) were identified using whole genome sequencing (WGS) data collected from six different datasets, representing over 20 distinct cancer types. The datasets included the Pan-cancer Analysis of Whole Genomes (PCAWG) study (n = 1950), the Hartwig Medical Foundation (HMF) project (n = 4823), the Personal Oncogenomics (POG) project (n = 570), The Cancer Genome Atlas (TCGA) (n = 724), the Clinical Proteomic Tumor Analysis Consortium (CPTAC) (n = 781), and the MMRF COMMPASS project (n = 758). Detailed data information, somatic variant calling methodology, and data processing are described in the Methods section (“WGS Mutation Data Collection and Processing,” Salvadores and Supek, 2024). Somatic variants were called on the hg19 reference genome. For the analysis of healthy tissues SNVs were obtained from single-cell-derived colonies from 4 different datasets(Lodato et al. 2015; Blokzijl et al. 2016; Brunner et al. 2019; Yoshida et al. 2020).

### Genomic data by gene type

The genomic coordinates of each tDNA were obtained using the UCSC LiftOver tool (https://genome.ucsc.edu/cgi-bin/hgLiftOver) to convert the previously obtained tRNA-scanSE 2.0 annotations from the T2T assembly to hg19. This approach utilizes the completeness of the T2T assembly, which is gapless and corrects previous uncertainty and scaffold mistakes, ensuring that the analysis benefits from the most accurate and comprehensive data available. A total of 537 tDNAs were identified in hg19 (Supp. Table. 3).

The data for other Pol III-transcribed genes (excluding tDNAs) was extracted from Pol3Base (Cai et al. 2022). Some genes in the database appeared in multiple experimental datasets, for that reason they were merged to maintain a set of unique genes. The Alu genes were also discarded. To convert the coordinates from hg38 to hg19 and to classify the genes by biotypes BioMart (https://www.ensembl.org/info/data/biomart/index.html) was used. Finally, pol III-transcribed genes used for the analysis were classified into specific biotypes: small nuclear RNA (snRNA), 5S ribosomal RNA (rRNA), and miscellaneous RNA or unclassified RNA (miscRNA) with a total of 106, 125, 80 genes, respectively.

### Mutation density

The analysis was performed using the hg19 reference genome. This procedure was used to analyze mutational density in cancerous samples for tDNAs, protein-coding genes, and other Pol III-transcribed genes (rRNA, snRNA, and miscRNA), as well as in non-cancerous samples for tDNAs. First, to ensure data quality, a set of quality filters was applied to the genomic data. To minimize errors due to misalignment of short reads and select uniquely mappable regions, reducing potential mapping ambiguities, all regions in the genome defined in the ’CRG Alignability 75’ track (Derrien et al. 2012) with alignability < 1.0 were masked out. In addition, regions that are unstable when converting between GRCh37 and GRCh38 (Ormond et al. 2021) were removed, as well as the ENCODE blacklist of problematic regions of the genome (Amemiya et al. 2019). Somatic SNVs were analyzed for different gene types: tDNAs, protein-coding genes, and other Pol III-transcribed genes, along with their flanking regions (e.g., 10 kb upstream and downstream, divided into 100-nt windows, with the gene itself designated as ’window 0’). For tDNAs and other Pol III-transcribed genes, only windows > 20 nucleotides that passed the specified quality filters were included, while for protein-coding genes, only windows > 100 nucleotides were considered. Only genes where window 0 fulfilled this threshold and with at least 10 mutations across all windows were included in the analysis. To normalize mutational densities by trinucleotide context (tri-nucleotide composition), we adjusted mutation counts based on the local sequence composition within each window. This involved calculating the frequency of each trinucleotide context’s occurrence and determining the mutation count for that context within each sample. Finally, we obtained the total mutation density for each window by adding the mutational densities across all trinucleotide contexts. The mutational density within each group (e.g., by transcription levels or gene type) was normalized to account for variations in the total mutation count across groups by dividing the observed mutation density in each window by the total mutation density within that group.

### Analysis for mutational density in tDNAs by tissue and age

To assess the relative mutation density at each tissue type, we calculated the fold change (FC) by dividing the mutation density of the tDNAs (window 0) to the average mutation density in sets of 10 consecutive windows in the flanking regions, (e.g., windows 1–10, 11–20, …, 91–100) both upstream and downstream. Only tissue types with a sample size greater than 20 (n > 20) were included in the analysis. Additionally, samples from the colorectum and uterus were stratified into subgroups based on previously described genomic instability patterns. These subgroups included microsatellite instability samples (MSI) or microsatellite stable (MSS) and hypermutated samples (HYPER) due to POLE deficiency. For age-related analysis we stratified the data by tissue, only tissues with a sample size greater than 200 (n > 200) were considered. To assess the fold changes by age the same approach was used to compare the mutational density between tDNAs and flanking. Spearman correlation was used to assess the relationship between mutation levels and age.

### Mutation frequencies in tDNAs at single base resolution

To analyze mutation patterns along tRNA sequences and identify mutation hotspots (specific nucleotides within tRNA sequences that accumulate mutations more frequently) the mutation frequency at each position in each tDNA was calculated. This was done by dividing the number of samples exhibiting mutations at a given position by the total number of samples analyzed. The set of tDNA positions that did not pass the quality filters, including alignability < 1.0, regions that are unstable when converting between GRCh37 and GRCh38, and ENCODE blacklist problematic regions, were removed from the analysis. For each tDNA the genomic coordinates were adjusted to align with the consensus tRNA reference positions (Beuning and Musier-Forsyth 1999; Biela et al. 2023), ensuring accurate nomenclature and allowing for the effective grouping of results across different tRNAs. To determine positions with significantly higher or lower mutation frequencies, *p*-values were calculated using the Wilcoxon test, comparing mutational frequencies for each position from all the tRNAs against all other positions, with adjustments for multiple comparisons using the Benjamini-Hochberg method. To assess the mutagenic load within the structural domains or regions within the tRNAs, a sliding window approach was applied. An 11-nucleotide window was moved along the tRNA sequence one nucleotide at a time, and the mutation frequencies within each window were combined to have a unique value by windows. The same statistical procedure used for the comparison by position was applied to calculate the differences by windows. In both analyses to consider a position or windows to be significant a threshold of adj. p-value < 0.01 and a mean higher or lower than the quantiles of 0.95 and 0.05 was considered. The mean mutational frequency at each position/windows across tDNAs was obtained and used to represent the results. To identify chimeric tRNAs, we first calculated the mutational density at the anticodon position (34, 35, 36) for tRNA^Ala^, tRNA^Leu^, tRNA^Ser^, and tRNA^Tyr^ by isodecoders. To detect specific mutations at the anticodon, absolute mutation counts and the annotation of base changes (e.g., C<G) were used to identify mutations occurring at the anticodon of tDNAs.

### Mutational signature

Skin cancer samples were excluded from the analysis due to their distinct mutation profiles. Specifically, skin cancers exhibit a high prevalence of mutations caused by ultraviolet (UV) light exposure. These UV-induced mutations create distinctive signatures that can overshadow or confound the detection of other mutational processes when analyzing diverse cancer types. To deconvolute the possible processes that cause mutations specifically at tRNAs, we applied the standard mutational signatures methodology(Supek and Lehner 2015; Alexandrov et al. 2020), with several modifications. First, as input instead of calculating the mutation frequencies across the 96 trinucleotides using WGS or WES, we calculated the tri-nucleotide mutation frequencies in subsets of specific sites of the genome, specifically in 4 groups of tRNA genes (divided by levels of gene expression) and their flanking regions (10 kb upstream and downstream, split in 4 chunks of 2.5 kb). As well as, 4 groups of protein coding genes (divided by levels of gene expression) and their flanking regions (10 kb upstream and downstream, split in 4 chunks of 2.5 kb). To account for confounders in the flanking regions, we removed 1 kb upstream of the tRNA (TSS) and all genes overlapping the flanking regions (i.e. protein coding exons and tRNAs).

Second, since we are studying a very small subset of the genome (e.g. tRNA genes), to gain statistical power we merged all mutations across different samples from the same cancer type, except samples that are MSI or POLE mutant that we kept as a different group. Additionally, we included as a separate group all somatic mutations from healthy samples. Third, because the genomic sites are different for each tri-nucleotide feature depending on the sample, for each sample and trinucleotide feature combination, we normalized the counts by the tri-nucleotide composition of each specific genomic subset.

In sum, the samples we considered for this analysis are a combination of tissue (cancer type or healthy), gene type (protein coding or tRNA), gene expression (levels 1 to 4) and position (tRNA, flanking upstream or downstream). To the matrix of normalized mutation frequencies across the 96 tri-nucleotides for these samples we applied the standard mutational signatures methodology (Supek and Lehner 2015; Alexandrov et al. 2020).

Specifically, we first applied bootstrap resampling (R function UPmultinomial from package sampling) to the normalized mutation frequencies. Next, we applied the non-negative matrix factorization (NMF) algorithm (R function nmf from package NMF) to the bootstrapped matrices, testing different values of the rank parameter (2 to 30), herein referred to as nFact. We repeated the bootstrapping and NMF 100 times for each nFact. We pooled all the results by nFact and performed a k-medoids clustering (R function pam from package cluster), with different number-of-clusters k values (2 to 30). We calculated the silhouette index value, a clustering quality score (which here measures, effectively, how reproducible are the NMF solutions across runs), for each clustering to select the best nFact and k values.

Based on the silhouette index (Second worst SI score) we selected the results from nFact=13 and k=13 (Supp. Fig. 3). From the selected option signatures with SI < 0.4 (Supp. Table. 4) were removed, and 12 signatures remained. These results include two matrices: H and W. The W matrix (our mutational signatures) describes the weight of each tri-nucleotide for every extracted signature. By comparing our signatures with the COSMIC database signatures (COSMIC_v3.3.1_SBS_GRCh37), we can match our signatures with the corresponding signature from COSMIC (Supp. Table. 5). Of note, we merged the two APOBEC signatures from COSMIC (SBS2+SBS13), since in our analyses they cannot be separated, likely because of our merging of samples. The H matrix describes the exposures or activities of each specific signature in every sample. Additionally, we stratified the exposures by tissue type and expression level to assess tissue-dependent and transcription-associated rate differences. To select relevant NMF-extracted signatures, different factors were examined. First, a cosine similarity threshold was applied to quantify the resemblance between NMF-extracted signatures and COSMIC signatures: cosine similarity values above 0.5 indicate moderate similarity, while values exceeding 0.85 denote strong similarity to COSMIC signatures. Additionally, we examined the signatures spectra (Supp. Fig. 5) to confirm their consistency with the spectra reported by COSMIC Furthermore, signature sample exposures and their relevance to specific cancer types were evaluated to ensure that the identified signatures are biologically meaningful. To identify signatures enriched in tDNAs, we calculated the fold change (FC) in the signatures exposure/activity levels between each gene type (tDNAs or protein-coding genes) and their respective flanking regions. To assess the abundance of hairpin loop structures in tDNA genes relative to the rest of the genome and to other RNA Polymerase III (Pol III) transcribed genes, we utilized a genome-wide dataset of predicted 3–5 nucleotide hairpin sites from Buisson et al. (https://www.science.org/doi/epdf/10.1126/science.aaw2872). Using the GenomicRanges package in R, we identified overlaps between hairpin regions and genomic coordinates of annotated tDNA genes, as well as with a separate dataset of other Pol III-transcribed genes. From this, we quantified the number of nucleotides within tDNA genes, other Pol III genes, and the remainder of the genome that were part of a predicted hairpin structure.

To calculate the frequencies of DNA hairpin structures in tDNAs that may he number of nucleotides not involved in hairpin structures was obtained by subtracting hairpin-associated nucleotides from the total nucleotide content of each corresponding genomic region (tDNA genes, other Pol III genes, and non-tDNA regions of the genome). To determine whether hairpin structures were statistically enriched or depleted in tDNA genes, we conducted two chi-square tests using the chisq.test() function in R: one comparing tDNA genes to the rest of the genome, and another comparing tDNA genes to other Pol III-transcribed genes. The number of nucleotides not involved in hairpin structures was calculated by subtracting hairpin-associated nucleotides from the total nucleotide content of each corresponding genomic region (tDNA genes, other Pol III genes, and non-tDNA regions of the genome). To determine whether hairpin structures were statistically enriched or depleted in tDNA genes, we conducted two chi-square tests using the chisq.test() function in R: one comparing tDNA genes to the rest of the genome, and another comparing tDNA genes to other Pol III-transcribed genes.

### Software and packages

The mutational analyses were performed using R v.4.1.2. Relevant R packages used were liftOver v.1.18.0, GenomicRanges v.1.46.1, Biostrings v.2.62.0, sampling v.2.10, matrixStats v.1.4.1, NMF v.0.26, ComplexHeatmap v.2.10.0, cluster v.2.1.2, dplyr v.1.1.4, tidyr v.1.3.0, ggplot2 v.3.5.1.

## Discussion

Proteome variability can have both negative and positive impacts upon cellular homeostasis. In microbes, for example, several adaptive mechanisms based on the induction of widespread proteome variation have been described, several of which are based on the use of mistranslating tRNAs (Ribas de Pouplana et al. 2014). In animals, adaptive proteome variations aided by changes in cellular tRNA populations are well known, and always linked to adaptive genetic switches that are important for both healthy tissues and tumor development (Pavon-Eternod et al. 2013; Gingold et al. 2014; Earnest-Noble et al. 2022). Adaptive mistranslation (the deliberate induction of translation errors during protein synthesis) has not been reported in animals, and it is assumed that such errors will always be deleterious to human health.

Frailty, sarcopenia, and many forms of neurodegeneration common during the aging process are linked to losses in proteome quality (Anisimova et al. 2018). Although such insults can ultimately be linked to ineffective and/or inaccurate gene translation, functional defects in the translation machinery are not generally considered significant contributors to aging (Anisimova et al. 2018). Similarly, human neoplasies that develop resistance to existing chemotherapies often do so through point mutations that thwart drug-recognition sites, but the possibility that such modifications arise through mistranslation events has not been explored (Roszkowska 2024).

We have studied in detail the dynamics of somatic mutagenesis in human tDNAs, a phenomenon previously observed in several model organisms and in human datasets (Saini et al. 2017; Sakhtemani et al. 2019; Sui et al. 2020; Seplyarskiy et al. 2023). Our data shows that human tRNA genes, but not other genes transcribed by Pol III, are hotspots of somatic mutagenesis that accumulate genetic changes directly as a function of their transcriptional rate. The fact that other Pol III-transcribed genes behave differently in terms of somatic mutagenesis indicates that some tDNA-specific factors drive these mutational events.

In agreement with previous studies, our mutational signature analysis indicates that members of the APOBEC3 family are the main contributors to this mutational load. Our topological analyses, together with published interactome data (Jang et al. 2024), indicate that the recognition of sequence motifs within tDNAs, coupled to interactions between APOBEC3 enzymes and tRNA modification enzymes, may be driving the specific mutational process at tDNAs (Supp. Fig. 8). Given that tRNA transcription and processing occur simultaneously, the specificity of somatic mutations at tDNA loci may be the result of specific interactions of APOBEC3 enzymes with components of the tRNA processing machinery in the vicinity of positive strands of tDNAs during transcription.

In tumors, but not in healthy cells, we find mutational signatures linked to chemotherapeutic agents used in cancer treatment. Interestingly these mutations display a clear anti-correlation with tDNA transcription levels. This could be due to the loss of tumor cells that incorporate such mutations in highly transcribed genes, and might indicate that chemotherapy induces a shift in the pattern of tDNA expression during tumor growth (Fig. 2e). Tumor samples display a similar behavior with regards to mutations caused by defects in PolE, but these mutations are again absent in healthy cells (Fig. 2e).

Analysis of the mutational load within tDNAs at single-nucleotide resolution shows that universally conserved positions in tRNAs display significantly reduced heterogeneity, indicating that these sites may be under negative (purifying) selection in somatic cells. Unexpectedly, we find anticodon positions to experience mutational loads no different from the average values in the whole sequence of tDNAs. This fact leads to the remarkable conclusion that somatic mutagenesis at actively transcribed tDNAs generate chimeric tRNAs with anticodon mutations previously shown to cause generalized substitutions in the human proteome. It stands to reason that age-related accumulation of mutations at tDNAs will result in an increase of such chimeric tRNAs in aging cells and tissues.

Our analysis of tDNA somatic mutations in different human tumors shows tumor-specific differences in the levels of mutation accumulation. Bladder cancer (BLCA), which displays the highest accumulation of such mutations, is characterized by highly augmented levels of APOBEC3A activity, and a several APOBEC-associated driver hotspot mutations have been identified in this malignancy (Shi et al. 2020; Lindskrog et al. 2021). This suggests that the high levels of tDNA mutations in BLCA are the consequence of the high levels of APOBEC activity in these tumors, and opens the possibility of a functional relationship between mutated tRNAs and BLCA development. Interestingly, brain cancer lays at the opposite spectrum in terms of tDNA somatic mutations, which are almost absent in the datasets from these tumors.

Whether tDNA mutations play a general role in carcinogenesis is an important question. This could happen, first, at the level of proteome variations that might compromise the activity of tumor suppressors or generate proteins that favor tumor growth, such as oncoproteins produced by translational variations in existing transcription factors. Secondly, mutant tRNAs with defective sequences or aberrant post-transcriptional modification patterns could alter the population of tRNA fragments (tRFs) in cells. It has been described that a dysregulation in the levels of specific tRFs is related with the aggressiveness of BLCA tumors (Papadimitriou et al. 2020). Finally, proteome variability induced by mutagenic tRNAs may provide tumoral cells with fitness advantages such as avoidance of immune recognition. Although such possibility has not been explored in human cells, in *Candida albicans* proteome variability induced by mistranslating tRNAs induces morphology changes that shield this organism against the immune response of its human host (Bezerra et al. 2013).

The extent to which tDNA somatic mutations accumulate in healthy tissues, their mutational signatures, and their physiological impact remain to be determined. However, the accumulation of defective tRNAs can rationally be linked to the many functional connections between tRNA dynamics and cancer (Zhang et al. 2018; Gupta et al. 2022). The age-dependent accumulation of mutations at tDNAs is apparent from the analysis of cancer datasets stratified by patient age, a factor that may directly impact upon the proteostasis of aging tissues (Schmidt and Schimmel 1993; Anisimova et al. 2018; Lopez-Otin et al. 2023). Inactive tRNAs in aging cells may induce an attenuation of protein synthesis efficiency and fidelity, either through their interference with components of the translation machinery. On the other hand, chimeric tRNAs would lead to mistranslation, possibly causing more acute problems such as protein aggregation, stress responses and inflammation, or the generation of aberrant antigenic peptides.

## Acknowledgements

We are grateful to Professors Manuel Santos (Multidisciplinary Institute of Aging, Porto), Francisco Real (CNIO, Madrid), Marc Martí-Renom (ICREA and CRG, Barcelona) and Nikolai Slavov (NorthWestern University, Boston) for helpful discussions. This work was supported by grants PID2023-147328OB-I00 and PID2019-108037RB-100 to L.R.d.P., and grants XXX to F.S.

